# Distinct neural variables underlie subjective reports of attention

**DOI:** 10.1101/2020.01.23.916841

**Authors:** Stephen Whitmarsh, Christophe Gitton, Veikko Jousmäki, Jérôme Sackur, Catherine Tallon-Baudry

## Abstract

Attention is subjectively experienced as a unified cognitive effort. This unitary experience of attention contrasts with the diversity of neural and peripheral correlates of attention. While the effects of attention on stimulus-evoked responses, alpha oscillations and pupil diameter, are well-known the relationship between these indices and the *subjective* experience of attention, has not yet been assessed. Participants performed a sustained (10 s to 30 s) attention task in which rare (10%) targets were detected within continuous tactile stimulation (16 Hz). Trials were followed by attention ratings on an 8-point Likert scale. Steady-state evoked fields (SSEFs) in response to tactile stimulation, as measured by magnetoencephalography, provided an objective measure of sensory processing. Beamformer source analysis of somatosensory alpha power was used as a measure of cortical excitability, while pupillometry provided a peripheral index of arousal. Attention ratings correlated negatively with contralateral somatosensory alpha power, and positively with pupil diameter. The effect of pupil diameter on attention ratings extended into the following trial, reflecting a sustained aspect of attention related to vigilance. The effect of alpha power did not carry over to the next trial, and furthermore mediated the effect of pupil diameter on attention ratings. Variation in SSEF power reflected stimulus processing under the influence of alpha oscillations, but were not readily expressed through subjective ratings of attention. Together, our results show that both alpha power and pupil diameter are reflected in the subjective experience of attention, albeit on different time spans, while continuous stimulus processing might not be metacognitively accessible.

**Significance Statement:** Attention is subjectively experienced as a unified cognitive effort, in contrast with a diversity of neural and peripheral measures shown to correlate with attention. We present the first comprehensive study on the complex inter-relationship between the most common bio-physiological indices of attention, and their association with the subjective experience of attention. We show that the subjective experience of attention correlates negatively with cortical alpha oscillations, and positively with pupil diameter. The latter reflected a sustained aspect of attention, spanning several tens of seconds, related to vigilance. Alpha power, representing cortical control, fluctuated faster and mediated the effect of pupil diameter on attention. Tactile steady-state power reflected stimulus processing, also under the influence of alpha oscillations, but did not contribute much to subjective ratings of attention.

## 2 Introduction

Subjectively, attention appears to be a clear and unified phenomenon (”Everyone knows what attention is” (James, 1890)), implicit in common parlance (”Pay attention to what she’s saying”), and our instructions to cognitive experiments (”Attend to the left of the screen”). Such a unitary concept of attention is in stark contrast with the diversity of neurophysiological measures that have been shown to correlate with attention, such as alpha oscillations, evoked responses to sensory stimulation, and pupillometry.

Alpha oscillations (≈ 10 Hz) reflect an attentional mechanism by which cortical excitability is modulated under cognitive control (Klimesch, Sauseng, & Hanslmayr, 2007; Jensen & Mazaheri, 2010). Increased somatosensory alpha has been shown to result in a reduced performance in tactile detection (Weisz et al., 2014) and discrimination (Haegens, Handel, & Jensen, 2011). Importantly, fluctuations in alpha power have been shown to correspond to retrospective subjective reports of attention (Macdonald, Mathan, & Yeung, 2011; Whitmarsh, Barendregt, Schoffelen, & Jensen, 2014; Whitmarsh, Oostenveld, Almeida, & Lundqvist, 2017). Attention has also been shown to increase the amplitude of steady-state evoked potentials/fields (SSEPs/SSEFs) (Giabbiconi, Trujillo-Barreto, Gruber, & Mller, 2007; Keitel, Andersen, & Mller, 2010; Keitel et al., 2019), with trial-by-trial variability corresponding to fMRI BOLD patterns associated with attentional control (Goltz et al., 2015). SSSEPs/SSEFs occur when sensory stimuli are repetitively delivered at a high enough rate that the relevant neuronal structures do not return to their resting states (Regan, 1989). They appear with the same fundamental frequency as that of the stimulus (Snyder, 1992; Tobimatsu, Zhang, & Kato, 1999), and have their origin in primary somatosensory cortices (Snyder, 1992; Tobimatsu et al., 1999). Finally, pupillometry provides an auxiliary index for attention, with both baseline (tonic) pupil diameter, as well as pupil responses, known to be reduced when attention is not on the task (Smallwood et al., 2011, 2012; Franklin, Broadway, Mrazek, Smallwood, & Schooler, 2013; Mittner et al., 2014; Konishi, Brown, Battaglini, & Smallwood, 2017; Kang, Huffer, & Wheatley, 2014).

Given these well-established correlations with attention, there has been surprisingly little investigation of their inter-relationship. Murphy et al. (2011) and Hong et al. (2014) found that pre-stimulus pupil diameter varied with N2 and P3 amplitude in EEG. Hong et al. (2014) found a negative quadratic relationship between between pre-stimulus pupil dilation and pre-stimulus alpha during an auditory oddball paradigm. However, this might be explained by an increase in *visual* alpha during *auditory* attention, limiting conclusions regarding the relationship between auditory alpha, pupil diameter and attention. Keitel et al. (2010, 2019) showed that attention modulates visual alpha oscillations and steady-state responses in opposite directions, but found no negative trial-by-trial correlations (Keil, Pomper, Feuerbach, & Senkowski, 2017).

Together, these results show that while several neurophysiological indices correlate with attention, it remains unclear how they A) relate to each other, B) are integrated into a unified subjective experience of attention. The current study is an attempt to address this issue by relating spontaneous fluctuations in neuronal oscillations, stimulus evoked fields, and pupil diameter, with subjective ratings of attention. We used a sustained attention task in which subjects detected rare (10%) target stimuli within trials of continuous tactile steady-state stimulation, followed by subjective attention ratings. We hypothesized that attention ratings would be positively correlated with pupil diameter and the amplitude of contralateral somatosensory steady-state responses, while negatively correlated with contralateral somatosensory alpha power. We furthermore tested whether these measures explain unique trial-by-trial variance, and explored their different temporal dynamics. Finally, we explored whether alpha power meditates the effect of arousal on attention ratings and cortical responses.

## 3 Materials and method

### 3.1 Subjects

26 healthy subjects enrolled after providing written informed consent and were paid in accordance with guideline of the local ethics committee. The experiment was in compliance with national legislation and the code of ethical principles (Declaration of Helsinki). One subject was excluded from the analysis due to an implant that would make subsequent MRI scanning unsafe. Three subjects were discarded due to excessive number of artifacts (described below), resulting in 22 subjects (11 females, *age* = 20.4–30.2, 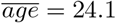, σ = 2.53).

### 3.2 Tactile stimulator

Stimulation was done at 16 Hz so to maximize the separation with the alpha frequency. The tactile stimulator consisted of a solenoid valve controlling air pressure transferred via a plastic tube to a tactor attached to the subject’s distal phalanx of their left index finger. Short opening of the valve resulted in the distention of a silicon membrane within the tactor. Air pressure, and the duration during which the valve was opened, were calibrated so that stimulation at resulted in a ‘tapping’ sensation rather than that of a continuous vibration. Delay between triggers and first moment of distension of the membrane was 33*ms*, for which all reported timings were corrected.

### 3.3 Experimental paradigm

The experiment consisted of a tactile detection task in subjects detected targets consisting of two missing stimulations within an otherwise continuous, rapid (16 Hz) stream of tactile stimulation (Figure 1). By studying attention in the tactile domain, pupillometry measurements remained free from visual confounds, and more clearly reflected changes in attention. Duration of the tactile stimulation was between 10 and 30*s* according to a truncated exponential distribution at a mean of 15*s*, approximating a flat hazard rate. Targets occurred on 10% of trials, at a random moment between after one second after onset, until one second before offset. An 8-step attention rating was presented one second after stimulation offset. Subjects were instructed to report their level of attention at the moment of stimulation offset. The attention rating was followed directly by a question on the presence or absence of the target. Responses were given by button-presses of the right hand.

**Figure 1:**
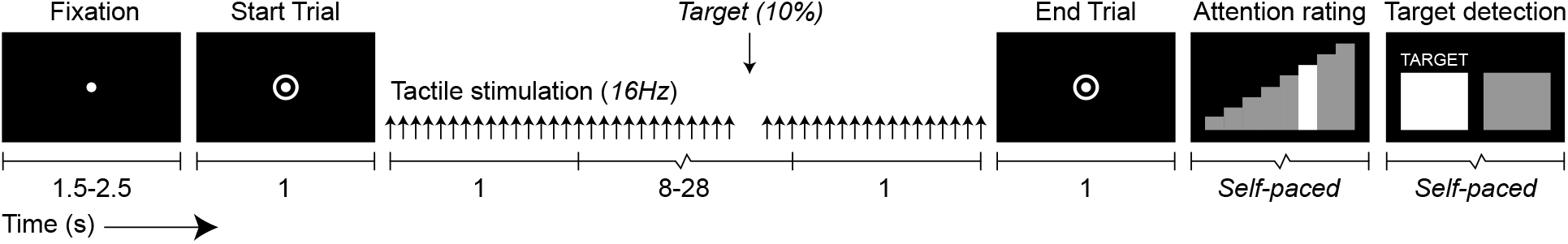
Subjects received tactile stimuli by mean of a pneumatic tactor attached to the distal phalange of the left index finger, and responded by means of button-presses with their right hand.

### 3.4 Procedure

After digitization of the head-shape and head-position coils, digitized with Polhemus Fasttrack (Polhemus Inc.), subjects were seated in the MEG with the tactor attached to the right hand. Subjects then completed a training session of ten trials in which half of the trial contained targets, to familiarize them with the procedure. Subjects then completed the experiment in four blocks of 50 trials, with a short break after the first and third block, and a longer break after the second block, in which they were lowered out of the MEG helmet. Care was taken to re-position subjects in the same position as the initial measurement, using in-house developed real-time head-localization of the MEG system.

### 3.5 Data acquisition

Horizontal eye-movements and eye-blinks were monitored using horizontal and vertical bipolar electrooculography (EOG) electrodes, placed on either side in line with the eyes, and above and below the right eye, respectively. Cardiac activity was monitored with bipolar electrocardiography (ECG) electrodes attached at the right clavicle and under the left lower rib. Impedance of electrodes was < 10*k*Ω measured with Siggi II impedance meter (Easycap GmbH). MEG measurements were carried out using a 306-channel whole-scalp neuromagnetometer system (Elekta Neuromag TRIUX^TM^, Elekta Oy, Helsinki, Finland) at the Centre de Neuroimagerie de Recherche (CENIR MEG-EEG) in Paris, France. Data was recorded at 1 kHz, low-pass filtered at 330 Hz and stored for off-line analyses. Eye position and pupil-diameter were monitored with an EyeLink 1000 (SR Research) and simultaneously recorded with the MEG, ECG, and EOG data. Anatomical MRI scans (Siemens, MPRAGE, 0.8cm isometric voxel size, GRAPPA=2) for source localization were either recorded at CENIR after MEG measurements, or acquired from participation in a previous study, recorded at the same location.

## 4 Analysis

### 4.1 Data selection

Target trials and false-alarm trials were discarded from further analysis. For each subject, trials were median-split between high and low-attention trials according to the individual distribution of attention-ratings. Trials were time-locked to the end of the stimulation, i.e. one second before the onset of the attentional probe. All comparisons between high and low attention were performed on the last 10 seconds, i.e. the minimum trial length, providing identical amount of data in each trial.

### 4.2 Artifact removal

Continuous MEG data were preprocessed off-line with MaxFilter 2.2.10 (Elekta Oy, Helsinki, Finland), using tSSS for artifact removal and head movement compensation with a correlation limit of ≥ 0.95 using a single data segment (Taulu & Kajola, 2005; Taulu & Simola, 2006). MEG data then analyzed using the Matlab-based Fieldtrip toolbox (Oostenveld, Fries, Maris, & Schoffelen, 2011) and custom functions. Data containing movement, muscle or superconducting quantum interference device (SQUID) jumps were detected by both automatic (based on z-value threshold) and visual inspection, and removed from further analysis. Trials wherein more than 20% of data contained artifacts were removed completely. Data was then decomposed into independent components using ICA (Makeig, Bell, Jung, & Sejnowski, 1996). Components reflecting EOG or ECG artifacts were iteratively removed if they correlated more that than three standard deviations (based on all remaining components channels) with either the EOG or ECG. Subjects of which the data had a combined percentage of artifacts that was larger than 3*σ* compared to all subjects, were rejected (3 rejected, 22 remaining), resulting in a subject-average of 12% artefacts (*σ* = 8%).

### 4.3 Alpha power sensor-level analysis

Alpha power (8 Hz to 14 Hz) was estimated on the 10-second trial segments. High versus low attention trials were compared using paired t-tests. Channels that showed a significant difference (*p* < 0.01) were included in a cluster-based permutation tests (Maris & Oostenveld, 2007) to identify spatial clusters of significant difference (*p* < 0.05, two-sided corrected, 4000 permutations).

### 4.4 Alpha power source reconstruction

Source reconstruction was done using a frequency-domain beamformer approach (Dynamic Imaging of Coherent Sources, DICS), which uses adaptive spatial filters to localize power in the entire brain (Gross et al., 2001; Liljestrm, Kujala, Jensen, & Salmelin, 2005). The brain volume of each individual subject was discretized to a grid with 5*mm* resolution. For every grid point, a spatial filter was constructed from the cross-spectral density matrix and lead field. Lead fields were calculated for a subject specific realistic single-shell model of the brain (Nolte, 2003), based on individual anatomical MRI images normalized to the International Consortium for Brain Mapping template (Mazziotta et al., 2001). The conductivity model of two subjects was based on a high-res template (Mazziotta et al., 2001) due to a lack of appropriate T1 MRIs. The spatial filter was based on all trials, for the whole stimulation period, to obtain an accurate and unbiased estimation. At each grid point and for each trial, alpha power at high and low attention trials were estimated on the last 10 second of each trial. Voxels that showed a significant relative difference 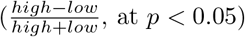 were used in cluster-based permutation tests (Maris & Oostenveld, 2007) to identify spatial clusters of significant difference (*p* < 0.05, two-sided corrected, 4000 permutations).

### 4.5 Steady-state source reconstruction

To determine the origin and attentional modulation of the steady-state response, data was band-pass filtered between 1 and 30*Hz*, segmented into intervals of 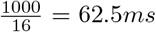 time-locked to the stimulation. Responses to each stimulation were then averaged within trials to obtain a trial-based ERP with a high signal-to-noise ratio. The first second of each trial was discarded, to allow the steady-state response to settle. We then determined the peak latency of the largest absolute evoked response over all sensors, and modeled the neural data of all sensors at this latency using a single dipole (Lutkenhoner, 1998), constrained to the same 5*mm* grid and using the same conduction model as for the beamformer procedure. In one subject, the latency of the largest absolute evoked response had to be adjusted by hand. Dipole locations were visualized using BrainNet Viewer (Xia, Wang, & He, 2013).

### 4.6 Evoked responses to stimulation onset and offset

The dipole solution for the steady-state response was used as an individual functional localizer for the primary somatosensory regions, and used to extract trial-by-trial evoked responses to stimulation onset and offset. Dipole solutions have an arbitrary polarity, because an inverse of the *signal* gives an equally good explanation of the data as an inverse of the *dipole* direction. To allow averaging over subjects, the orientation of the evoked responses were therefor matched through an iterative procedure: the polarity of the subject’s averaged evoked response was flipped if this resulted in a higher correlation with the average evoked response over subjects. Peak latencies of the evoked components were then determined based on the average over both high and low attention ratings and all subjects, after which subject-level amplitudes were compared between attention levels using paired t-tests.

### 4.7 Steady-state power analysis

Time-locked responses to periodic steady-state stimulation retain the phase of stimulation, allowing signal-to-noise to be improved by averaging the time-courses over trials before power estimation. The power of the 16*Hz* steady-state response was therefore determined on the averaged time-course over trials, time-locked to the last stimulation. A Fast-Fourier transform was calculated over the last 10 seconds. The power estimates of 16*Hz* responses during the last 10 seconds were then compared for high versus low attention trials using a paired t-test.

### 4.8 Pupillometry

Pupillometry data was first band-pass filtered between 2 and 150*Hz* and time-locked to the last stimulation. Saccades and artifacts were detected by means of z-thresholding (*z* > 2), padded by 150*ms* and linearly interpolated. Trials containing more than 50% of unusable data were removed. Differences between high and low attention were normalized to relative change and compared using cluster-based permutation tests (Maris & Oostenveld, 2007) to identify temporal clusters of significant difference (*p* < 0.05, two-sided, 4000 permutations).

### 4.9 Relationships between physiological signals

Models of the relationships between physiological measures were tested using mixed-effects linear models (Bates, Mchler, Bolker, & Walker, 2015; Kuznetsova, Brockhoff, & Christensen, 2015), with subject as a random factor. Different models were compared by means of a Bayes Factor, derived from Bayes Information Criteria, as follows (Wagenmakers, 2007): *BF*_01_ ≈ exp(Δ*BIC*_10_/2). Mediation analysis were performed using the mediation toolbox (Tingley, Yamamoto, Hirose, Keele, & Imai, 2014), using bootstrapping (*n* = 1000) for testing direct and indirect (mediated) effects.

## 5 Results

### 5.1 Task performance

Subjects were able to perform the task accurately, with 99% (*σ* = 0.01%) correct rejections of non-target trials, and 87% (*σ* = 0.1%) correct detection of targets. Attention ratings showed a supra-normal distribution, with a mode of 6, indicating that subjects were generally able to maintain attention (Figure 2A). The influence of trial duration, trial number (within a block) and block number (1 to 4), as well as the interaction between trial number and block number were modelled together in a mixed effects linear model, with subject as random effects. The association between trial length and attention rating did not appear to be linear (Figure 2B), and was not found to be significant (*t*(3055) = −0.71, *p* = 0.78). Time-on-trial, as modelled as the interaction between trial number and block number, was also not of influence on attention rations (*t*(3051) = −0.71, *p* = 0.78). However, trial number was found to be a significant predictor (*t*(3051) = −4.404, *p* =< 0.001), as, on average, each block started with high attention ratings, which stabilized after ≈ 10 trials (Figure 2C).

**Figure 2:**
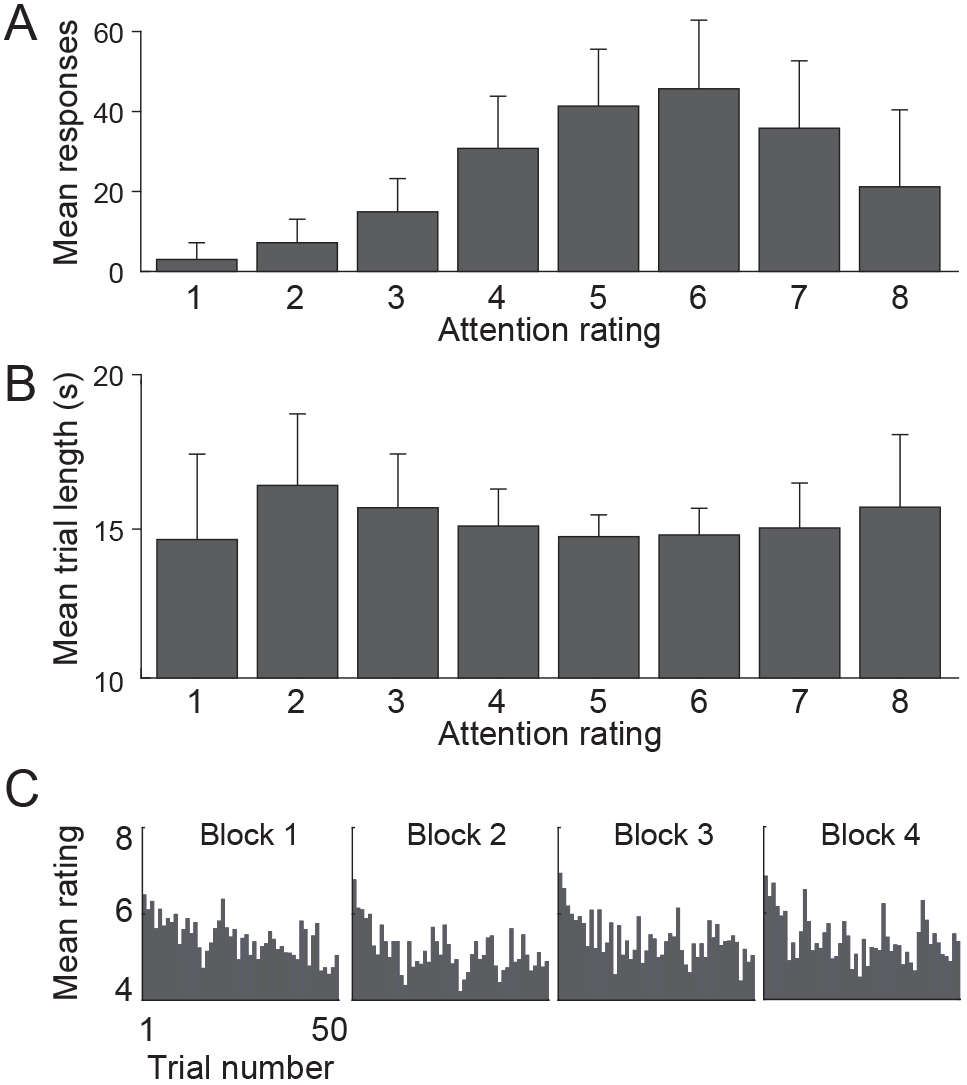
Overview of attention rating responses (1 = lowest, 8 = highest). A) Average attention ratings over participants show a mode of 6. B) Trial-durations averaged over participants did not differ between ratings. C) Subjects started on average with higher attention early on in each block.

### 5.2 Somatosensory alpha reflects self-reported attention

We tested whether alpha power during stimulation co-varied with attentional rating. We indeed found that higher attention ratings corresponded to reduced alpha power at right central sensors, contralateral to the attended hand (Figure 3A, *p*^*cluster*^ = 0.0165). Source-level analysis showed the source of the contralateral alpha suppression to be at the primary somatosensory region (Figure 3B & Table 1).

**Table 1:** Anatomical labels for clusters of significant effect of attention on alpha power, for all regions *>* 30 voxels. Four clusters were identified: *p* = cluster probability (Monte Carlo), Peak *t* = maximum t-value in cluster, Mean t = average t-value in cluster, Size = number of voxels, % = percentage activation of anatomical region, X, Y & Z = Talairach coordinates.

| Cluster | $p$ | Region (AAL) | Peak $t$ | Mean $t$ | Size | % | X | Y | Z |
| --- | --- | --- | --- | --- | --- | --- | --- | --- | --- |
| 1 | 0.0012 | Supramarginal Gyrus (R) | -4.35 | -4.06 | 188 | 9.5 | 42 | -38 | 40 |
|  |  | Inferior Parietal Lobule (R) | -4.32 | -3.98 | 144 | 10.7 | 40 | -40 | 40 |
|  |  | 'Postcentral Gyrus (R) | -4.09 | -3.90 | 61 | 1.6 | 46 | -32 | 46 |
| 2 | 0.0012 | 'Cerebellum Crus2 (L) | -4.91 | -4.15 | 47 | 2.5 | -20 | -88 | -32 |
|  |  | 'Cerebellum Crus1 (L) | -4.98 | -4.10 | 93 | 3.6 | -18 | -84 | -30 |
| 3 | 0.0040 | 'Angular Gyrus (R) | -4.17 | -3.95 | 42 | 2.4 | 62 | -50 | 34 |
|  |  | 'SupraMarginal Gyrus (R) | -4.06 | -3.92 | 26 | 1.3 | 62 | -48 | 34 |
| 4 | 0.0080 | 'Heschl Gyrus (R) | -4.00 | -3.89 | 29 | 11.6 | 46 | -22 | 10 |

**Figure 3:**
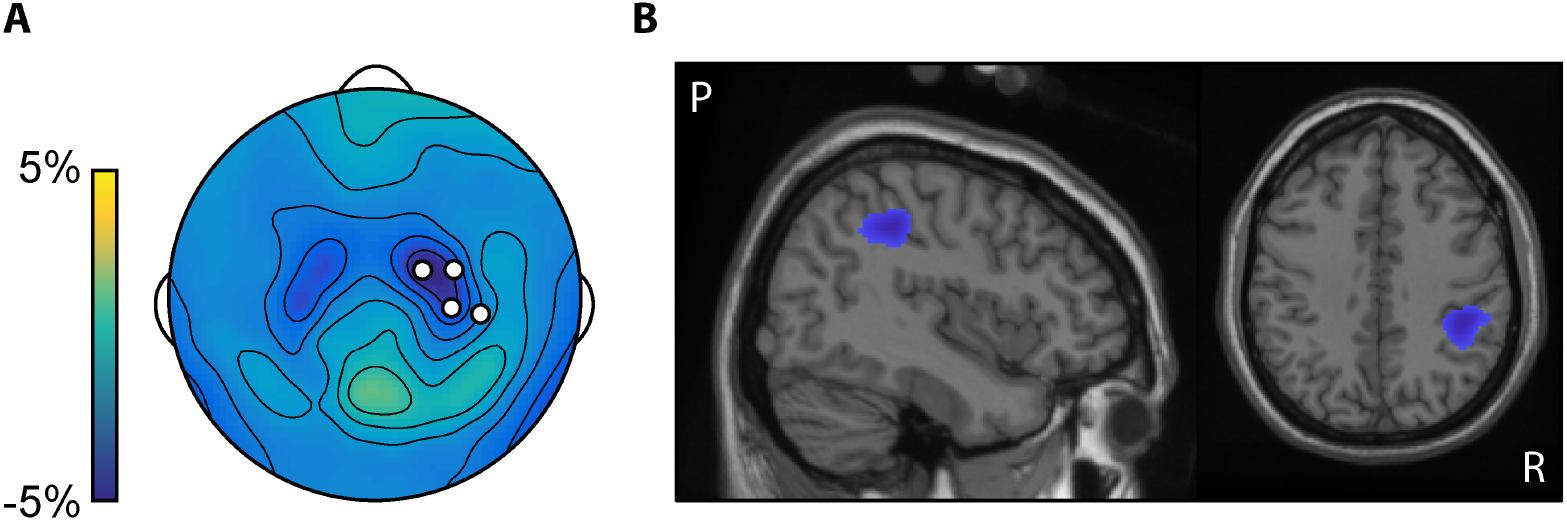
A) Topography of differences in 8 Hz to 14 Hz power on sensor-level, channels of significant cluster indicated in white. B) Beamformer localization showed a significant cluster in alpha power at somatosensory cortex, with a maximum at *X* = 42.0 mm, *Y* = −38.0 mm, *Z* = 40.0 mm

### 5.3 Tactile evoked and steady-state responses reflect self-reported attention

We then tested whether the tactile stimulation resulted in a steady-state response at stimulation frequency. Spectral analysis of the average SSEF indeed showed a clear peak at 16*Hz* with a contralateral central topography (Figure 4A) that was maintained until the end of the trial (Figure 4C). A comparison of 16*Hz* power, averaged over the last ten seconds, showed a significant increase in high over low attention trials (*t*(21) = 3.33, *p* = 0.0032, Fig 4B). To verify the somatosensory origin of the steady-state response, we performed a dipole localization on the evoked response averaged over stimulations. We found that SSEF dipoles were indeed located at the right primary somatosensory cortex (Figure 4D). The onset of the steady-state stimulation evoked a contralateral somatosensory evoked field at 50*ms* and 85*ms* (Figure 5A). Peak amplitudes after steady-state stimulation onset did not differentiate between high and low attention (*t*(21) = 0.80, *p* = 0.43, and *t*(21) = −0.39, *p* = 0.70, respectively). The evoked field stabilized within 500*ms* into a clear steady-state response at stimulation frequency (Figure 4E). Stimulation offset (Figure 4E) also evoked a response at 48.5*ms*, 136.5*ms* and 209.5*ms*, with significantly larger deflections for high-attention trials only in the late component (*t*(21) = −1.09, *p* = 0.28, *t*(21) = 1.79, and *p* = 0.087, *t*(21) = −2.13, *p* = 0.045, respectively), however, these findings would not survive a correction for multiple comparisons for testing three components.

**Figure 4:**
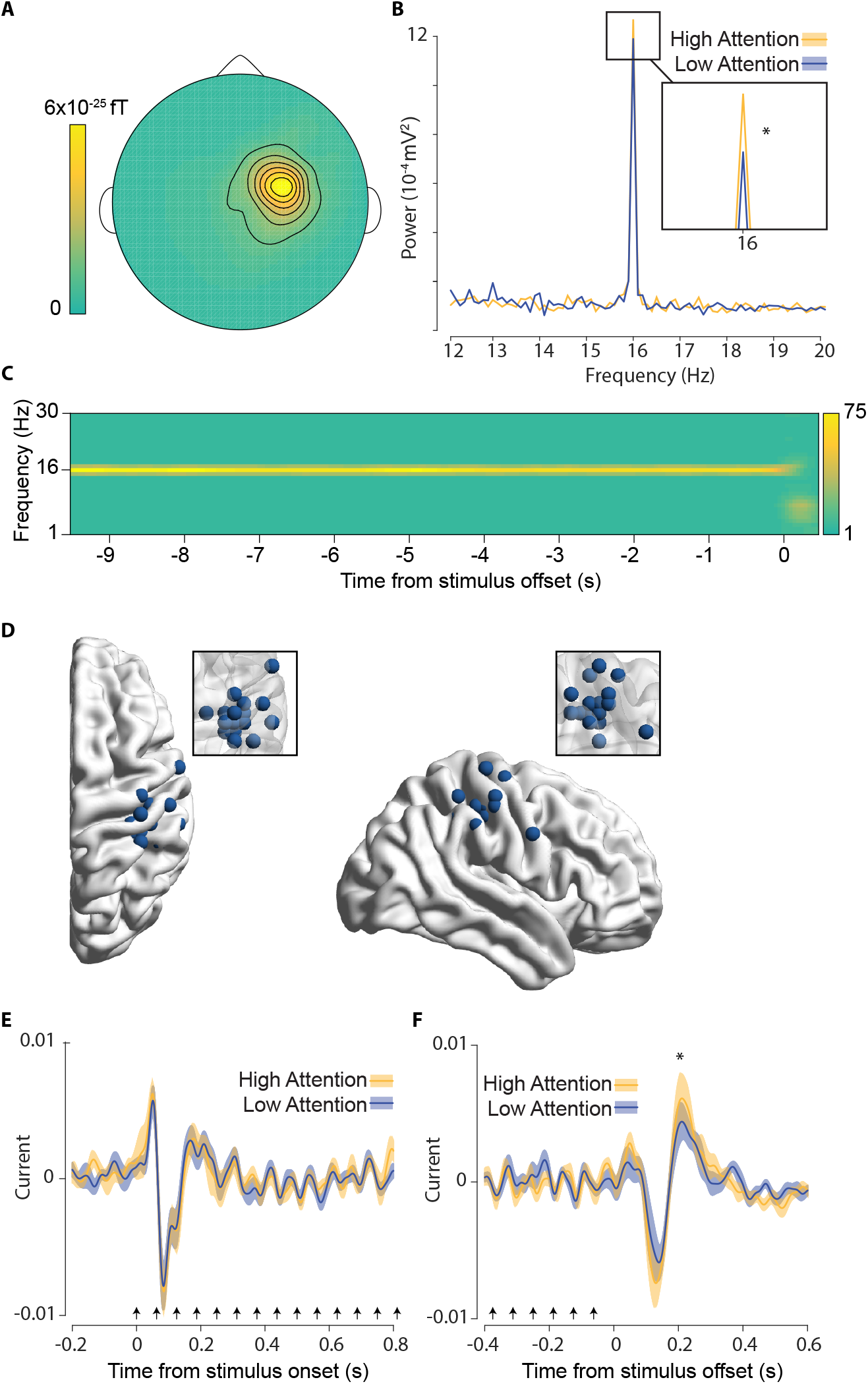
A) Topography of 16 Hz power. B) High attention trials showed higher SSEF power during trials. C) Baseline-corrected 16Hz power, showing maintenance of 16 Hz SSEF throughout the trial, up to 75x power compared to post-stimulus baseline. D) Dipole localization of steady-state response for each subject. E) Source reconstructed ERF time-locked to trial onset. F) Source reconstructed ERF time-locked to trial offset. Arrows in B & C indicate steady-state stimulation

**Figure 5:**
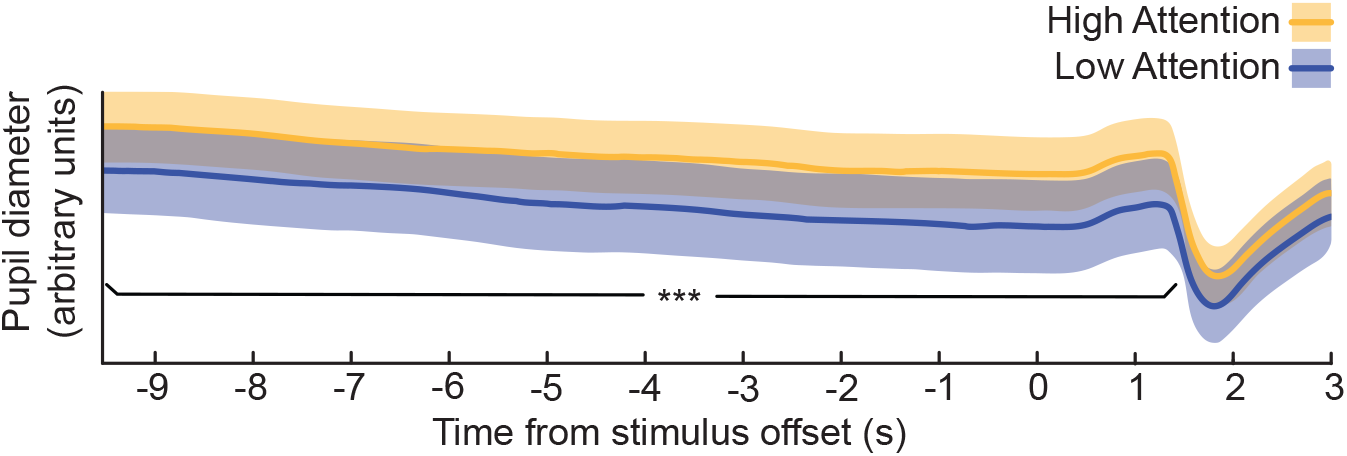
Pupil diameter over time for high and low attention ratings showing significantly larger pupil diameter during high attention trails.

### 5.4 Pupil-size reflects self-reported attention

To test whether pupil diameter also co-varied with attentional rating, we performed a cluster permutation test of pupil diameter. Pupil diameter was indeed found to be larger for trials in which subjects reported to be more attentive. This effect was sustained in time, with a single temporal cluster extending from 10 seconds before stimulation offset until 1.68*s* after (Figure 3; *p*^*cluster*^ = 0.006).

### 5.5 Alpha and pupil explain unique variance in attention ratings

Our results show that pupil diameter, contralateral somatosensory alpha power, and steady-state responses, distinguish between high and low attentional ratings. To investigate whether pupil-diameter, contralateral alpha power and steady-state power explain unique rather than shared variance in explaining attention ratings, we modeled their trial-by-trial fluctuations as fixed factors in a mixed model of attention ratings with subject as a random factor. To take into account the effect on attention ratings of time on task, we included both *block number* (1 to 4) and *time-within-block* as confound regressors. Values for alpha power, pupil diameter, and steady-state power were all taken from their individual analysis, i.e. based on ROI or dipole location over the last ten seconds of stimulation in each trial, and z-transformed to simplify interpretation and comparability of the model estimates.

Alpha power (*t*(3070) = −6.1, *p* = 1.16 ×10^−9^) and pupil diameter (*t*(3070) = 11.271, *p* < 2 ×10^−16^) both explained unique variance in predicting attentional ratings, indicating that they reflect different underlying components of the (self-reported) attentional state (Table 2). However, steady-state 16Hz power did not contribute to attentional ratings in this trial-by-trial analysis (*t*(3070) = −0.541, *p* = 0.59). A model-comparison between a model *with* steady-state power and a model *without*, clearly indicated that the model *without* steady-state power explained attention ratings better (*BF* = 31.5). We therefore removed steady-state power from further analysis of alpha power and pupil diameter. The lack of an effect of steady-state responses stands in contrast with the results from the median-split analysis. The steady-state response might therefore be redundant when modelling both alpha and pupil measures. However, in the median-split analysis the phase-consistency of the steady-state response was exploited by averaging the time-courses *before* the frequency analysis, greatly reducing variations in the signal that were not phase-locked to the stimulation.

**Table 2:**
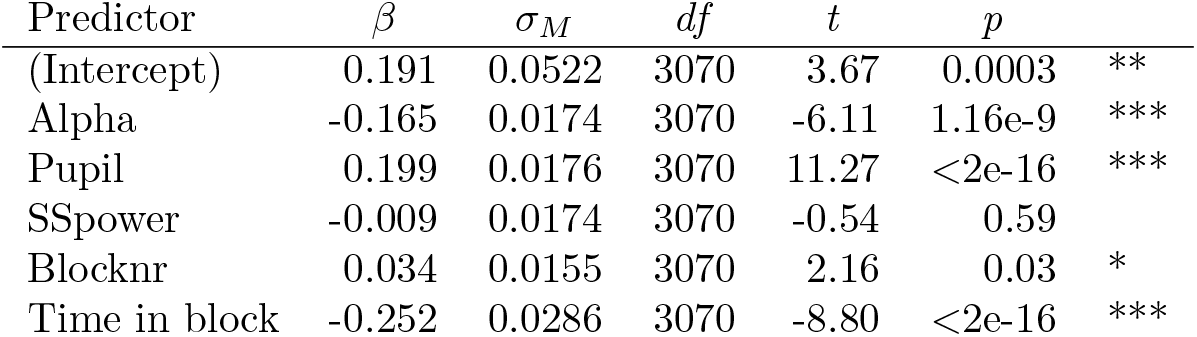
Mixed model of trial-by-trial attention ratings

| Predictor | $\beta$ | $\sigma_M$ | $df$ | $t$ | $p$ | |
| --- | --- | --- | --- | --- | --- | --- |
| (Intercept) | 0.191 | 0.0522 | 3070 | 3.67 | 0.0003 | ** |
| Alpha | -0.165 | 0.0174 | 3070 | -6.11 | 1.16e-9 | *** |
| Pupil | 0.199 | 0.0176 | 3070 | 11.27 | <2e-16 | *** |
| SSpower | -0.009 | 0.0174 | 3070 | -0.54 | 0.59 |  |
| Blocknr | 0.034 | 0.0155 | 3070 | 2.16 | 0.03 | * |
| Time in block | -0.252 | 0.0286 | 3070 | -8.80 | <2e-16 | *** |

### 5.6 Pupil diameter tracks tonic state of arousal

Pupil diameter is typically associated with a relatively tonic state of arousal, while alpha power is known to be fast-reacting to cognitive control. To test the different temporal effects of these measures on attention, we added trial-shifted predictors to our model. Specifically, we tested whether values of attention ratings, alpha power and pupil diameter of the *current* trial could explain attention ratings, alpha power and pupil diameter of the *next* trial. Indeed, a model including current and next trial values greatly improved the fit (Table 3, *BF* = 6.17 × 10^12^). Specifically, attention ratings at each trial were shown not to be independent from each other, but highly predictive of subsequent attention ratings (*t*(3068) = 9.05, *p* = 2 × 10^−16^). This shows that self-reported measures of attention persisted over trials. Interestingly, while pupil diameter predicted subsequent attentional ratings (*t*(3068) = −2.91, *p* = 0.004), alpha power did not (*t*(3068) = 0.006, *p* = 0.75). These results indicate that pupil diameter reflected aspects of attention that varied more slowly over trials, while alpha power reflected more transient trial-by-trial variations in attention.

**Table 3:** Mixed model of trial-by-trial attention ratings including previous trial (’).

| Predictor | $\beta$ | $\sigma_M$ | $df$ | $t$ | $p$ | |
| --- | --- | --- | --- | --- | --- | --- |
| (Intercept) | 0.170 | 0.052 | 3068 | 3.29 | 0.001 | ** |
| Alpha | -0.106 | 0.018 | 3068 | -6.04 | <1.74e-9 | *** |
| Pupil | 0.211 | 0.020 | 3068 | 10.70 | <2e-16 | *** |
| Rating' | 0.161 | 0.018 | 3068 | 9.05 | <2e-16 | *** |
| Alpha' | 0.006 | 0.018 | 3068 | 0.316 | 0.75 |  |
| Pupil' | -0.058 | 0.020 | 3068 | -2.91 | 0.004 | ** |
| Blocknr | 0.021 | 0.015 | 3068 | 1.34 | 0.18 |  |
| Time in block | -0.230 | 0.029 | 3068 | -7.90 | <3.74e-15 | *** |

### 5.7 Alpha power mediates effect of pupil on attention

Because pupil diameter was shown to influence attention ratings beyond the current trial, while alpha power explained attention ratings in a trial-by-trial manner, we explored whether the effect of pupil diameter on attention was also mediated by alpha power. Indeed, a mediation model (pupil → alpha → attention rating) showed that pupil diameter’s effect on attention was significantly mediated by alpha power (*β* = 0.008, *p* < 2 × 10^−16^). Statistical estimates of the predictors are reported in Figure 6A.

**Figure 6:**
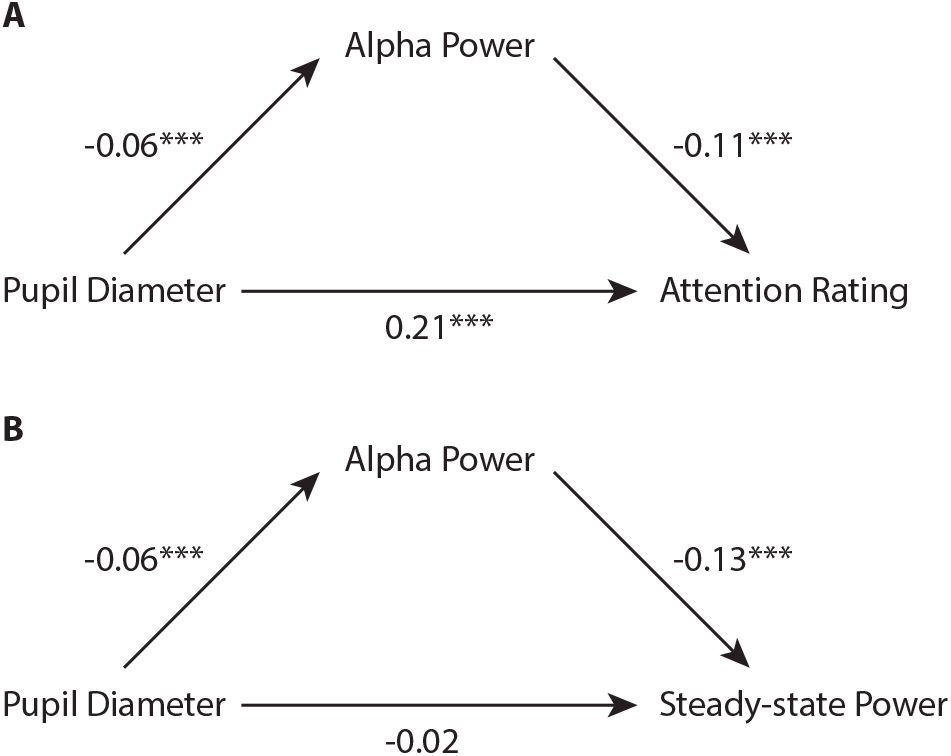
Schematic diagram of the direct and mediation effects of (A) pupil diameter on attention ratings meditated by alpha power, and (B) pupil diameter on steady-state power mediated by alpha power (B). Numbers indicate *β* values and corresponding significance (***: *p* < 0.001)

### 5.8 Exploration of steady-state power as alpha-dependent measure of attention

As reported above, steady-state responses did not explain attention ratings in a trial-by-trial analysis, when including alpha power and pupil diameter, but only when using a median-split approach. We therefor explored an alternative hypothesis that steady-state responses corresponds to aspects of attention that are better captured by pupil diameter or contra-lateral alpha power, rather than by self-reports. A mixed effects model indeed showed that alpha power (*t*(3068) = 6.48, *p* < 1.04 × 10^−^10) but not pupil diameter (*t*(3068) = −0.30, *p* = 0.76) explained steady-state power (Table 4). Interestingly, while no direct effect of pupil diameter on steady-state power was found, pupil diameter did influence steady-state power when mediated by somatosensory alpha power (figure 6B, pupil → alpha → steady-state: *β* = −0.008, *p* < 2 × 10^−16^). Together, these results suggest that steady-state power might not reflect the *subjective* experience of attention, but rather reflects attentional modulation of stimulus processing by alpha power.

**Table 4:** Steady-state power: Fixed effects in mixed trial-by-trial model including previous trial (’).

| Predictor | $\beta$ | $\sigma_M$ | $df$ | $t$ | $p$ | |
| --- | --- | --- | --- | --- | --- | --- |
| (Intercept) | 0.141 | 0.054 | 3068 | 2.60 | 0.009 | ** |
| Pupil | -0.006 | 0.021 | 3068 | -0.30 | 0.76 |  |
| Alpha | 0.120 | 0.018 | 3068 | 6.48 | <1.04e-10 | *** |
| Blocknr | -0.019 | 0.016 | 3068 | -1.19 | 0.24 |  |
| Time in block | -0.096 | 0.030 | 3068 | -3.25 | 0.001 | *** |
| Steady-state' | 0.043 | 0.018 | 3068 | 2.46 | 0.014 | * |
| Pupil' | -0.016 | 0.021 | 3068 | -0.76 | 0.45 |  |
| Alpha' | 0.033 | 0.019 | 3068 | 1.79 | 0.074 |  |

## 6 Discussion

The multifaceted nature of attention was clearly reflected by our findings that each of the three variables we studied (steady-state stimulation responses, endogenous fluctuations in pupil-diameter, and alpha power), relate to self-reported attention in unique ways. Endogenous fluctuations in contralateral alpha power, as well as tonic pupil diameter, strongly reflected the subjective attentional state. Furthermore, they did so according to different temporal dynamics: while alpha activity explained attention ratings on a trial-by-trial basis, pupil diameter showed an influence on attention ratings that also extended beyond the current trial. This is in line with the idea that pupil diameter reflects features of attention related to arousal such as motivation and vigilance, while alpha activity reflects momentary cognitive control. In fact, it is generally accepted that while alpha suppression can be under fast-acting cortical (attentional) control, pupil diameter is thought to be modulated subcortically, beyond conscious cognitive control. Furthermore, alpha power and pupil diameter did not act entirely independently: mediation analysis showed that the relationship between pupil diameter and the subjective experience of attention was in part mediated by alpha activity. This further reinforces the idea that cognitive control happens ‘on top of’ more tonic fluctuations of vigilance and motivation.

In contrast to alpha power and pupil diameter, steady-state responses corresponded only weakly to self-reported attention ratings in a median-split analysis, and this relationship disappeared in a trial-by-trial analysis when alpha power and pupil diameter were included as predictors. However, alpha activity was shown to have a strong effect on steady-state power, and pupil diameter influenced steady-state power (only) when mediated by alpha power. This suggests that while early somatosensory processing, as reflected by steady-state power, are under the influence of attentional processes (Eimer & Forster, 2003), this influence does not translate in large differences in the experience of attention. In other words, unlike the attentional processes associated with alpha power and pupil diameter, i.e. cognitive control and vigilance, respectively, stimulus processing as measured with SSEFs might not be metacognitively accessible. These findings are also in line with the idea that alpha power and steady-state responses represent different attentional effects, in which steady-state responses predominantly reflect an early cortical mechanism for tracking fluctuations in stimulus-specific visual input (Keil et al., 2017), while alpha suppression gates the access of sensory information to further downstream sensory-processing stages (Jensen & Mazaheri, 2010; Zumer, Scheeringa, Schoffelen, Norris, & Jensen, 2014). Only the latter might allow access to higher-order cognitive processes required for conscious awareness. It could be argued that the lack of an association between steady-state responses and attention ratings in the trial-by-trial analyses could be due to the fact that such an analysis does not benefit from the improvement of signal-to-noise due to averaging phase-locked responses, as was done is the median split analysis. However, the strong effect of alpha power on steady-state responses suggests that the signal-to-noise was sufficient.

Steady-state stimulation resulted in an unequivocal 16*Hz* contralateral somatosensory response lasting throughout the trial. The evoked response to stimulation onset at 50*ms*, corresponds to the early contralateral P50 (Hamalainen, Kekoni, Sams, Reinikainen, & Ntnen, 1990) derived from excitatory inputs in the upper cortical layers of area 3b (McLaughlin & Kelly, 1993). The offset response at 209.5*ms* was somewhat earlier than what has been reported for vibration offset responses (240*ms*, (Nangini, Ross, Tam, & Graham, 2006)), but can be explained by the known delay in response to vibration versus tactile stimulation (Hari, 1980; Johnson, Jrgens, & Kornhuber, 1980; Pfurtscheller, Schwarz, & Gravenstein, 1985).

As expected, contralateral somatosensory alpha power was shown to negatively correlate with subjective attention (Whitmarsh et al., 2017, 2014). While there has been much evidence that contralateral alpha suppression reflects *cued* spatial somatosensory attention (Klimesch et al., 2007; Jensen & Mazaheri, 2010; Jones et al., 2010; Haegens, Osipova, Oostenveld, & Jensen, 2010; Anderson & Ding, 2011; van Ede, Kster, & Maris, 2012; van Ede, Szebnyi, & Maris, 2014; Haegens et al., 2011), these results provide further evidence for the role of *spontaneous* (i.e. endogenous) fluctuations of alpha power in *sustained* spatial attention (Macdonald et al., 2011; Whitmarsh et al., 2014, 2017). We also found, as expected, that pupil diameter positively correlated with attention ratings. This adds to the growing evidence that pupil-diameter tracks fluctuations of sustained attention, as in mind-wandering (Smallwood et al., 2011, 2012; Franklin et al., 2013; Mittner et al., 2014; Konishi et al., 2017; Kang et al., 2014).

This study provides a first attempt to address the commutability between subjective and multiple objective measures of attention. Taken together, our results suggest that the subjective experience of attention is the result of an integration of different central and peripheral sources, each providing relatively independent information on the attentional state. These findings extend previous studies that show that spontaneous fluctuations of attention are metacognitively accessible (Macdonald et al., 2011; Whitmarsh et al., 2014, 2017), by showing that subjective reports of attention also explain variations in arousal (pupillometry). Our results also indicate a unique role of alpha oscillations in both subjective and objective measurements of attention, by mediating of the effect of arousal (pupillometry) on both sensory processing (SSEF amplitude) and attention ratings, and providing an association between sensory processing and the subjective experience of attention. In other words, attentional modulation of stimulus processing might not be metacognitively accessible in the subjective experience of attention. Instead, associations between attentional modulation of stimulus processing and the subjective experience of attention could be driven by an association with cognitive control, indexed by alpha power.

To conclude, our results show that not only “everyone knows what attention is” (James, 1890), but that our subjective experience of attention corresponds reflects its multidimensional nature. Future studies will need to investigate whether the experience of attention can also be *subjectively* dissociated, by probing participants subjectively on dimensions of sensory experiences, focused attention and arousal.

## 7 Acknowledgements

We would like to thank Margaux Romand-Monnier and Clemence Almeras for their help in subject planning and recording. This work was supported by funding from the European Research Council (ERC) under the European Unions Horizon 2020 research and innovation program (grant agreement No 670325, Advanced grant BRAVIUS) and senior fellowship of the Canadian Institute for Advance Research (CIFAR) program in Brain, Mind and Consciousness to C.T.-B., as well as from ANR-10-LABX-0087 IEC, ANR-10-IDEX-0001-02 PSL.

## 8 Contact information

Stephen Whitmarsh (corresponding author)

Phone: +33 (0)7 681 600 79

## Notes

Conflict of interest statement: The authors declare no competing financial interests.

#### Summary of Updates

Submitted version

